# Attention-Based Solution for Synergistic Virus Combination Therapy

**DOI:** 10.1101/2025.04.22.649915

**Authors:** Shayan Majidifar, Mohsen Hooshmand

## Abstract

Computational drug repurposing is vital in drug discovery research because it significantly reduces both the cost and time involved in the drug development process. Additionally, combination therapy—using more than one drug for treatment—can enhance efficacy and minimize the side effects associated with individual drugs. However, there is currently limited research focused on computational approaches to combination therapy for viral diseases. This paper proposes AI-based models to predict novel drug combinations that can synergistically treat viral diseases. To achieve this, we have compiled a comprehensive dataset containing information on viruses, drug compounds, and their approved interactions. We introduce two attention-based models and compare their performance with traditional machine learning and deep learning models in predicting synergistic drug pairs for treating viral diseases. Among all the methods tested, the random forest algorithm and one of the attention-based models utilizing a customized dot product as a predictor showed the highest performance. Notably, two predicted combinations—acyclovir + ribavirin and acyclovir + Pranobex Inosine—have been experimentally validated to produce a synergistic antiviral effect against the herpes simplex virus type 1, as reported in existing literature.

## Introduction

Viruses are among the oldest biological entities on Earth, having existed for billions of years [1]. They play essential roles in ecosystems, such as controlling the populations of organisms like bacteria and archaea [2], returning nutrients to ecosystems by breaking down plankton [3], and driving evolutionary processes [4]. However, viruses are often associated with diseases and have caused significant morbidity and mortality in recent decades. Within just two decades, three deadly single-stranded viruses—SARS-CoV, MERS-CoV, and SARS-CoV-2—have caused severe acute respiratory infections, leading to a substantial number of deaths regionally and globally [5]. Monkeypox, a double-stranded virus from the smallpox family, can cause symptoms such as rash, fever, and swollen lymph nodes [6, 7]. Additionally, HMPV (human metapneumovirus), a negative-sense single-stranded RNA virus from the Pneumoviridae family, causes respiratory tract infections in children, adults, and the elderly [8].

All of these viruses, among many others, emerge unexpectedly and often cause deadly diseases without any prescribed treatments readily available. Therefore, discovering effective drugs to stop or hinder these viral diseases is of utmost importance. On the other hand, drug discovery is a time-consuming process and an extremely expensive industry [9]. Relying solely on proposing new drugs in emergencies can endanger the lives of thousands or even millions of people. Thus, drug repurposing can be an effective solution in such scenarios. However, this approach also requires significant effort [10].

Using computational methods for screening and proposing approved drugs for repurposing is an efficient and effective strategy that significantly reduces costs and time [11]. Drug repurposing, especially with the advent of deep learning and transformer-based methods, has had a high impact on drug research and discovery. Most studies focus on monotherapy, where drug target predictions typically suggest single drugs for treating diseases. These targets can include proteins, viruses, and other biomolecules.

However, in recent years, there has been growing interest in predicting combination therapies for curing diseases [12]. The goal is to increase treatment effectiveness while simultaneously reducing side effects compared to higher doses of a single drug. Cancer treatment and oncology, in particular, have seen extensive investigations into synergistic combination therapies using machine learning and deep learning methods in recent years [13–16].

In contrast, combination therapies for treating viral diseases remain rare [17, 18]. This is due to challenges such as drug resistance, the difficulty of treating such viruses, and the lack of knowledge during emergency situations [19]. As a result, proposing synergistic antiviral combination therapies suffers from a lack of sufficient research. This work focuses on attention-based methods for predicting drug pairs to treat viral diseases.

The proposed approach for viral combination therapy in this work utilizes an attention mechanism for generating a significant embedding for the prediction of the interaction more effectively and efficiently. We prepared the attention-based models in two ways, i.e., prediction using an MLP and prediction using a customized inner product. The former is called the **V**irus **C**ombination **T**herapy using **at**tention and **MLP**, or VCTatMLP and the latter is called **V**irus **C**ombination **T**herapy using **at**tention and **Dot** product, or VCTatDot. Figure 1 and Figure 2 show two approaches, respectively. We organized the paper as follows. Section Related work reviews previous studies in this field. Section Basic definitions and methods outlines key concepts used in our approach. Section Dataset describes the dataset employed in this study. Section Proposed methods presents two approaches for virus combination therapy. Section Results reports the evaluation metrics and experimental findings. Finally, Section Conclusion summarizes the work and its contributions.

**Fig 1.**
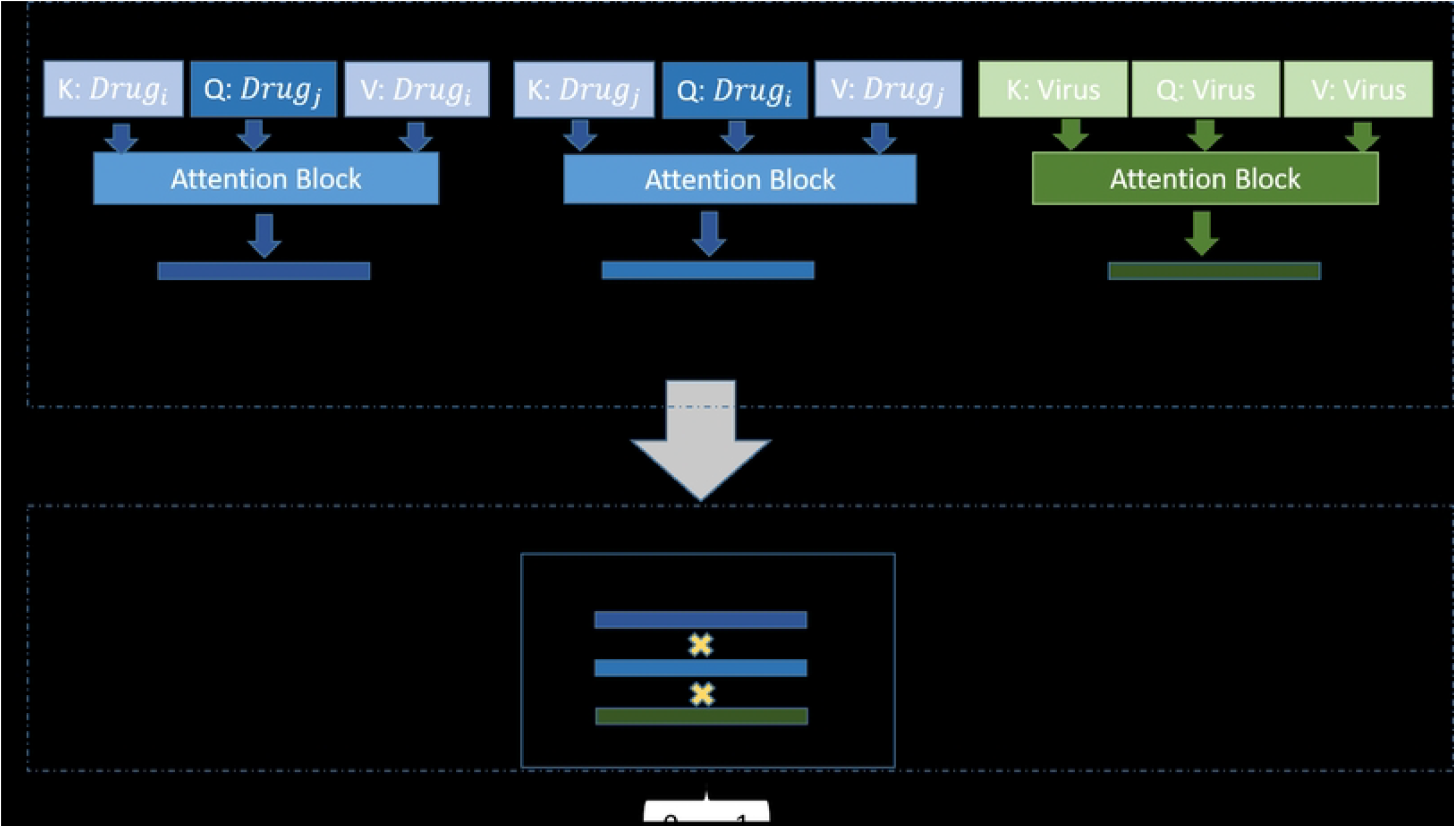
VCTatMLP architecture, the encoder section consists of attention mechanism blocks and the prediction section consists of an MLP block applied on the concatenation of the embedding vectors.

**Fig 2.**
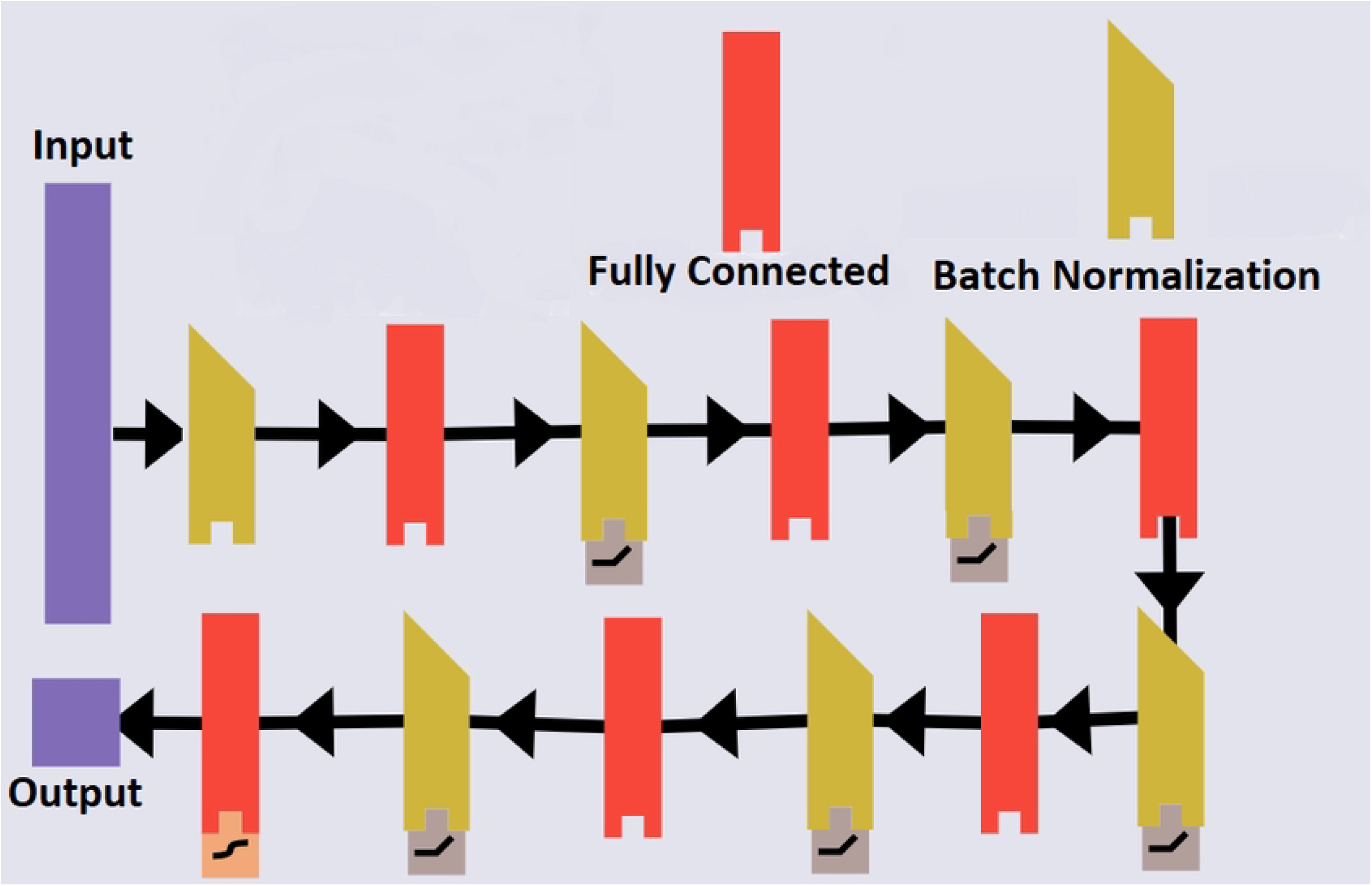
VCTatDot architecture, the encoder section consists of attention mechanism blocks and the prediction section consists of applying a sigmoid function on the tri-dot product of the embedding vectors.

## Related work

El-Hafeez et al [20] developed a framework for classifying and recommending synergistic drug combinations. They utilized the O’Neil drug interaction dataset [21], which contains 22737 drug combinations tested against 39 cancer cell lines. To classify the data into three categories—synergistic, additive, or antagonistic—they applied various classification methods, including naïve Bayes, random forest, k-Nearest Neighbors, and Logistic Regression. For the regression task, they employed linear regression, ridge regression, and random forest regression to predict combination sensitivity scores (CSS). Additionally, they enhanced model performance through data preprocessing by identifying outliers using the Interquartile Range (IQR) method and addressing data imbalance through random undersampling.

Jiang et al. [22] developed a graph convolutional network (GCN) for predicting synergistic drug combinations. Their approach utilized multiple biological interaction networks as input data. Specifically, they took advantage of three types of networks: drug-drug synergy (DDS), drug-target interaction (DTI), and protein-protein interaction (PPI) to construct their heterogeneous multi-type graph using graph embedding techniques. They formulated the problem as a link prediction task and trained their model using the O’Neil dataset. Their model was compared against traditional machine learning models, including random forest, support vector machine, elastic net, and deep neural networks, demonstrating superior performance. Furthermore, the GCN model successfully predicted synergistic drug combinations for various cancer cell lines, such as breast, colon, lung, and skin cancer, with some predictions later validated in clinical trials.

Yuan et al. [23] proposed a machine learning framework to model cellular responses to perturbations, including drug combinations. Unlike traditional black-box machine learning models, CellBox combines explicit mathematical models of molecular interactions with machine-learning-based parameter inference. Using ordinary differential equations (ODEs), CellBox learns molecular interaction networks from perturbation data and can predict responses to unseen drug combinations. The model was trained using reverse-phase protein array (RPPA) data, which includes measurements of protein and phosphoprotein levels taken before and after perturbation. In addition, the model was tested on a dataset of melanoma cell lines treated with various drugs. The study demonstrated that CellBox could accurately predict dynamic cellular responses to novel drug combinations, achieving an average Pearson correlation of 0.93 in cross-validation experiments. Furthermore, the inferred interaction networks showed significant overlap with known biological pathways, indicating that CellBox not only predicts drug synergy but also provides valuable mechanistic insights.

Güvenç et al. [24] provided a comprehensive review of machine learning approaches for drug combination therapies, organizing them into three main categories: drug combination sensitivity prediction, drug synergy prediction, and drug synergy classification. They highlighted that integrating diverse data sources enhances drug synergy prediction. However, challenges such as data standardization and missing values persist in this field of research.

The drug combination sensitivity prediction category focuses on determining how specific drug combinations impact cell viability or growth. Various machine learning (ML) models have been applied, including deep learning (DNNs), matrix factorization methods, and regression models. For instance, the BestComboScore [25] model utilizes DNNs to integrate multiple types of molecular and drug features. In contrast, the ComboFM [26] model employs factorization machines to learn higher-order drug interactions. The drug synergy prediction category quantifies the interaction strength between pairs of drugs. Güvenç et al. emphasized that the most effective models combine multi-source data, such as gene expression profiles, chemical structures, and protein-protein interactions. They discussed the DrugComboRanker [27] model, which uses Bayesian non-negative matrix factorization to cluster functionally similar drug pairs and predict their likelihood of synergy. Other studies have employed matrix completion techniques, like DECREASE [28], which reconstructs dose-response matrices to estimate synergy scores.

The last category is drug synergy classification, which involves classifying drug combinations as synergistic, additive, or antagonistic interactions. Classification models, including random forest, SVMs, and deep learning architectures, have been widely adopted. SyDRa [29], a random forest-based approach, integrates drug–chemical structure, drug–target network and pharmacogenomics data to identify effective drug combinations.

Virus combination therapy receives limited attention due to the narrow-spectrum effectiveness of antiviral treatments and the persistent challenge of drug resistance [19]. Computational and AI methods have been explored for COVID-19 combination therapy suggestions [17, 30]. Myhre [31] compiled a list of virus therapy drug combinations reported in various experiments (e.g., *in vitro, in vivo*, and clinical trials). Using Myhre’s list, previous research established an initial dataset for virus synergistic combination therapy [32] and employed machine and deep learning models to predict novel combination therapy candidates. New transformer and attention-based methods have been largely overlooked in predicting virus combination therapy. This work proposes attention-based methods to predict novel synergistic drug combinations for virus treatment.

## Basic definitions and methods

This paper proposes two attention-based methods for predicting synergistic effects of pair drugs for viral disease treatments and compares them with a machine learning algorithm consisting of a support vector machine, and random fores,t and a convolutional deep model (DRaW). We use several concepts for machine and deep learning, therefore, it is necessary to introduce the fundamental concepts. This section introduces and reviews the important concepts for the proceeding of the paper.

### Support Vector Machine

Support Vector Machine (SVM) [33] is a machine learning algorithm introduced by Vapnik, which has been utilized in this research. The algorithm aims to find optimal separating boundaries that maximize the margin between two different classes. However, in many cases, the two classes are not linearly separable. To address this issue, various techniques, such as applying different kernel functions and allowing some misclassification, are used to improve classification performance. A kernel is a function that maps input data into a higher-dimensional feature space, making it possible to transform non-linearly separable data into a space where it may become linearly separable. We employed three types of kernels: the linear kernel, which is suitable for linearly separable data and performs well when a straight-line boundary can be defined; the polynomial kernel, which is effective for data that follows a polynomial pattern with a specific degree; and the radial basis function (RBF) kernel, which is particularly useful for complex and nonlinear data structures. Another important aspect of SVM is the C parameter, which controls the balance between correct classification and misclassification of the model. The objective of SVM is to determine an optimal hyperplane that separates the data, but some misclassification may occur based on the dataset. A higher value of C makes the model more sensitive to classification errors, forcing it to correctly classify as many data points as possible, which may lead to overfitting. Conversely, a lower value of C allows for more misclassification, reducing sensitivity to errors, which can result in underfitting [34].

### Random Forest

Random Forest [35] is another machine learning algorithm used in this research. It belongs to the ensemble learning category, where instead of training a single classifier, multiple classifiers are trained, and the final label is determined based on majority voting. In the case of Random Forest, multiple decision trees are trained, each selecting a subset of features randomly from the available feature set. Each tree learns a model independently, and the aggregation of these trees helps improve the overall performance while reducing overfitting. To evaluate the importance of features at each decision tree level, two criteria were used: Gini and Log Loss. Each decision tree selects its own set of features based on these metrics. The Gini criterion measures how well a feature separates different classes; the lower the Gini value, the better the feature is at distinguishing between classes. It essentially evaluates the entropy reduction at each split, favoring features that contribute more to classification purity. The Log Loss criterion, on the other hand, assesses the model’s ability to predict the probability of each sample belonging to a given class. This criterion is highly sensitive to classification errors, making it useful for selecting features with better classification capability.

Another important parameter in Random Forest is tWhich has the following propertiehse max features, which determines the number of features used to train each decision tree. In this paper, we employed two selection strategies: log *n* and 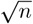, where n represents the total number of available features. The feature vectors in our dataset capture similarity measures for the drugs and viruses involved in each combination. Each combination consists of two drugs and one virus, resulting in a total feature vector length calculated as **d**_*i*_ + **d**_*j*_ + **v**_*k*_, where **d**_*i*_ and **d**_*j*_ indicate the number of drug-related features, and **v**_*k*_ represents the number of virus-related features. This structured approach ensures that each tree is trained with a diverse subset of features, enhancing the model’s robustness and generalization capabilities [36].

### Multilayer perceptron

These networks have been inspired by the structure and function of the brain. The brain contains billions of connected neurons and processes the information [37, 38]. Multilayer perceptron, similarly, uses the concept of neurons in an ordered manner. The MLP consists of several layers, each layer consisting of several perceptrons. The perceptrons of each two consecutive layers are connected, therefore, the output of a perceptron of the current layer is the input of the perceptrons of the next layer. Thus, each perceptron receives many inputs equal to the number of perceptrons from the predecessor layer multiplied by the corresponding weights. The MLP will learn these weights automatically using a large enough labeled dataset. To learn the nonlinear relation between the feature vector and the label, each layer applies an activation function on its output and then passes the results to the perceptrons of the next layer. This action takes place through the network. The learning and correction of the weights happen based on the comparison of the true labels and their corresponding predicted values. Using the error, the correction is back-propagated through the networks and updates the weights to avoid the errors. This network repeats this action to reduce the errors for all labels and reach convergence in the error minimization function at the end [39].

### Convolutional neural network

Convolutional neural network (CNN) is a type of deep network that was proposed for image processing tasks at first. Later, it was used in different areas of learning [40]. CNNs are composed of different layers, such as convolution layers, pooling layers, and fully connected layers. The convolutional layer is responsible for extracting essential patterns needed for accurate label prediction. In a convolutional neural network (CNN), this layer identifies patterns that range from simple to complex. Convolutional layers learn filters, or kernels, to extract these patterns, facilitating the prediction of results. The pooling layer uses a single representative of each block of the values, therefore reducing the volume of the data. The fully connected layer or MLP finds the relation among the found patterns in the data [41].

This paper uses a special type of CNN for the prediction of virus combination therapy. It is defined first at the [42, 43], and a customized version of that is used for combination therapy [32]. Therefore, it is a proper candidate for comparing the performance of proposed methods.

### Attention mechanism

The attention mechanism is a computational process that identifies and focuses on the important parts of the input data for higher-performance learning. Its main idea is based on assigning different weights to different parts of the input vector regarding the importance of each part automatically [44]. This leads to recognizing the dependencies of the far parts of the data. The attention mechanism is based on three main concepts **query, key**, and **value**, which we describe shortly. The *query* is a vector that the model looks for to find some information based on that. The *key* is a vector representing the information available in each element of the input. This vector is compared with the query vector. In other words, the similarity among keys and queries is calculated according to the following inner product of key and query.

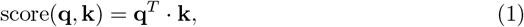

where **q** and **k** are *n ×* 1 vectors and Eq. (1) represents the similarity score computed based on inner product.

In the next step, the computed scores of all keys are converted to the interval [0, 1] using *softmax*. These outputs are known as attention weights, shown in the following equation.

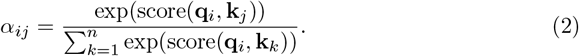

Where *α*_*ij*_ represents the similarity score between query vector **q**_*i*_ and the key vector **k**_*j*_. The final step of the attention mechanism involves summing the weighted value vectors as follows.

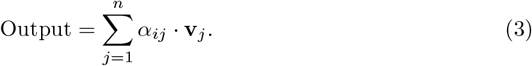

Where each input value vector **v**_*j*_ is multiplied by its corresponding attention weight *α*_*ij*_ and summed to obtain the overall weighted values. The whole procedure is compacted in the following equation.

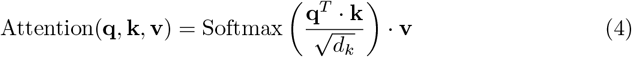

These attention weights *α*_*ij*_ are learned during the training phase. This paradigm, due to giving higher weights and attention to the more important parts of the inputs for outcome prediction has improved the performance of deep learning enormously.

### Self- vs. cross-attention

It is worth noting that the attention mechanism has two categories of self-attention and cross-attention [45]. The former considers all three vectors of query, key, and value the same; that is, each input vector is fed to the model as key, query, and value. In other words, this type of mechanism concentrates on the impact or the importance of the different parts of the same vector. The latter is another attention mechanism in which the query, key, and value vectors are not the same. For example, it accepts a sentence as a key and a value and its corresponding translation to another language as the query. In this work, we consider one drug as the query and another drug as the key and value, and vice versa.

### Tri-dot product

The inner product or dot product on a vector space 𝕍 ∈ ℝ^*n*^ is a function as,

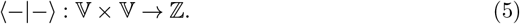

Which has the following properties [46].

1. ∀**v** ∈ 𝕍 : ⟨**v, v**⟩ ≥ 0. ⟨**v, v**⟩ = 0 iff **v** = **0**.
2. ∀**v**_1_, **v**_2_, **v**_3_ ∈ 𝕍:
  - ⟨**v**_1_ + **v**_2_, **v**_3_⟩ = ⟨**v**_1_ + **v**_3_⟩ + ⟨**v**_2_, **v**_3_⟩.
  - ⟨**v**_1_, **v**_2_ + **v**_3_⟩ = ⟨**v**_1_ + **v**_2_⟩ + ⟨**v**_1_, **v**_3_⟩.
3. ∀*c* ∈ ℤ & ∀**v**_1_, **v**_2_ ∈ 𝕍: ⟨*c*.**v**_1_, **v**_2_⟩ = ⟨**v**_1_, *c*.**v**_2_⟩ = *c ×*⟨**v**_1_, **v**_2_⟩.
4. ∀**v**_1_, **v**_2_, **v**_3_ ∈ 𝕍: ⟨**v**_1_, **v**_2_⟩ = ⟨**v**_2_, **v**_1_⟩.

In this paper, the vectors receive values from a subspace of ℝ^*n*^. Therefore, the inner product of two vectors **v** and **w** is,

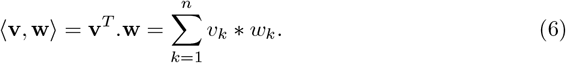

Where, *v*_*k*_ and *w*_*k*_ are the *k*-th values of **v** and **w**, respectively. The inner product has many applications in different areas of research and studies. For example, they are used for similarity comparison in quantum mechanics and computing [47]. Furthermore, it can be used in decoding or prediction of values in machine and deep learning approach [48]. This paper utilizes an inner product for the prediction of synergistic combination therapy for viruses. However, it uses a special type of that which is the generalization of three vectors instead of two vectors. The inner product of three vectors is called *genralized dot product* or *tri-dot product* [49], which is defined as follows.

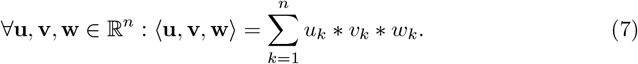

Where, *u*_*k*_, *v*_*k*_, and *w*_*k*_ are the *k*-th values of **u, v**, and **w**, respectively. Tri-dot product has the last three properties of the dot product, but it does not follow the first properties and can accept negative values for some of the vectors.

As mentioned above, due to generating a significant embedding vector by attention approaches, the decoding (prediction) step can be a tri-product. The results show its high performance which we mention in Section Results

## Dataset

This section provides information on the dataset utilized in this paper. Any deep learning model needs a suitable dataset to learn from and infer new predictions [50]. The starting point of this work is a dataset that contains information on the combination of viruses and drugs used for combination therapy. Myhre collected a list of combination therapies used for emerging infectious diseases from the literature. He showed that developing new tools for combating ineffective viruses plays an important role in disease treatment. He correctly mentioned that the use of more than one drug can lead to higher performance in the treatment of diseases rather than a single drug, so it is necessary to introduce combination therapy as an effective approach to handle viral diseases. In his list, the synergism effect of pair drugs on viruses has been collected from approved clinical trials, *in vitro*,and *in vivo*. However, the list does not follow a proper standard for drugs’ names and viruses. Therefore, we updated the list with the standard names of drugs and viruses regarding DrugBank and NCBI. Then, we generated a dataset from the information in the revised list of viral combination therapies. The drug SMILES [51] was collected from the DrugBank [52], and the nucleotide sequences of viruses were collected from NCBI [53]. Using Tanimoto score and sequence alignment, we computed the similarity among drugs and viruses, respectively. Then, for each drug, we produced a feature vector using its similarity values with other drugs. The same happens with viruses. Therefore, the data set contains the drug characteristic vectors *D*_211*×*211_, in which the *i*-th row (**d**_*i*_) represents the characteristic vector of the *i*-th drug. Furthermore, a virus feature vector has been generated using similar values between viruses called *V*_44*×*44_, where the *j*-th row **v**_*j*_ shows the virus feature vector *j*-th virus. This dataset is called CombTVir [32]. The drug-virus interactions are considered as labels in this dataset. The dataset contains 44 viruses, 211 drugs, and 372 approved drug-virus interactions. Therefore, it has a high percentage of sparsity.

## Proposed methods

The proposed methods in this paper, VCTatMLP and VCTatDot, are based on the attention mechanism, illustrated in Figure 1 and Figure 2. Both methods consist of two steps, that is, encoding and decoding. The former generates a significant embedding vector of each input entity. Encoding is an attention-based approach to generating embeddings, and its general procedure has been defined in Section Attention mechanism. The encoding is identical for both proposals. The decoding predicts the synergistic effect of drugs on a given virus. We propose two methods for the prediction of combination therapy for viruses. The two proposed methods differ in the decoding step.

VCTatMLP is the first model, which consists of two sections: encoder (embedding generation for viruses and drugs) and decoder (prediction of the drug-drug-virus interaction), shown in Figure 1. It aims to predict the synergism effect of combination drug i and drug j on virus k, where 1 *≤ i, j ≤ n* and *i* ≠ *j* and 1 *≤ k ≤ m*. It utlizes three attention blocks for the encoding section. For drug embeddings, the first two attention blocks are utilized for their embedding generation. The first attention block uses drug i as the key and value, drug *j* as the query, and its output is the embedding for drug *i*. The next attention block for the drugs is drug j as the key and value and drug *i* as the query. The two mentioned attention blocks belong to the cross-attention category. Therefore, the cross-relation among two drugs leads to a higher significant embedding for each drug. Lastly, for viruses, it uses the third attention block, a self-attention block, in which its *q, k*, and *v* vectors are set to virus input vectors. Therefore, three embeddings are generated for drug i, drug j, and virus k, called **d**_*i*_, **d**_*j*_, and **v**_*k*_, respectively. Notably, the encoding section for both proposed methods is identical; therefore, VCTatDot follows the same line of g embeddings for drugs and viruses.

The significance of the encoder section is that we can use a lighter version of methods for decoding or prediction. VCTatMLP utilizes an MLP for the goal of prediction. Figure 3 shows the MLP architecture used for the prediction. It contains one batch normalization layer as the first layer, five fully connected layers, each followed by a batch normalization layer, and a final fully connected layer for outputting the prediction value. The activation function of the inner FC layers is ReLU, and the last activation layer is sigmoid.

**Fig 3.**
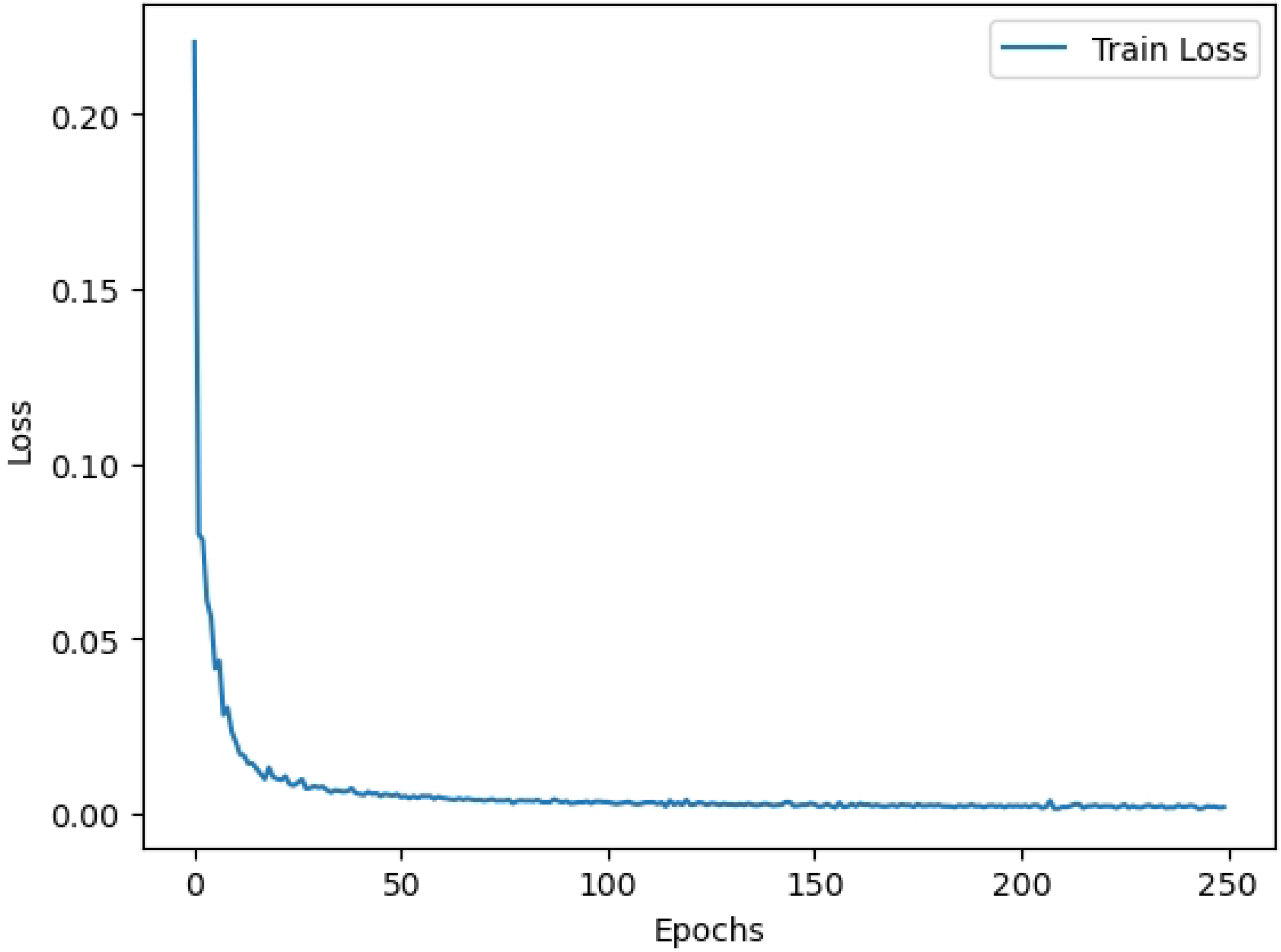
Architecture of MLP section of VCTatMLP

VCTatDot utilizes a customized version of the inner product for synergism prediction, tri-dot product, introduced in Section Tri-dot product-Eq. (7). By having embeddings **d**_*i*_, **d**_*j*_, and **v**_*k*_, the following equation computes the synergism prediction.

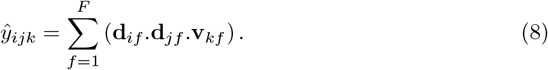

Here, *F* is the embedding dimension, and **d**_*if*_, **d**_*jf*_, and **d**_*kf*_ represent the *f* -th elements of embeddings **d**_*i*_, **d**_*j*_, and **d**_*k*_, respectively. As shown in Eq. (8), VCTatDot calculates the element-wise product of these three embeddings and sums the results. Briefly, both methods employ the proposed attention mechanism to generate significant representations of two drugs and the virus. VCTatMLP and VCTatDot utilize MLP and tri-dot product, respectively, for downstream tasks, both using *binary cross-entropy* as the loss function.

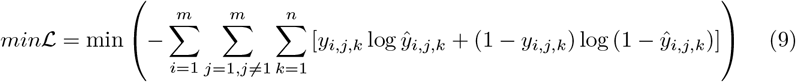

## Results

The proposed methods were implemented in Python using scikit-learn for machine learning and PyTorch for deep learning models. Experiments were conducted on a system running Ubuntu 22.04 LTS with an Intel Xeon E5 v4 (4 cores), 16 GB RAM, and an RTX 3060 12G GPU. Table 1 details the specific libraries and versions used.

**Table 1.** Library/Framework used for developing the proposed methods.

| Library/Framework | Version |
| --- | --- |
| **numpy** | **1.24.3** |
| **matplotlib** | **3.4.2** |
| **pandas** | **1.5.2** |
| **rdkit** | **2024.3.3** |
| **sklearn** | **1.2.1** |
| **torch** | **2.0.1 + cu118** |

We evaluate algorithm performance using Accuracy, MCC, AUC-RoC, and AUPR, calculated from the true positives (TP), true negatives (TN), false positives (FP), and false negatives (FN) of the confusion matrix, using the following equations.

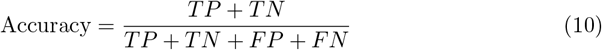

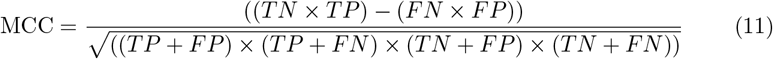

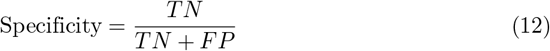

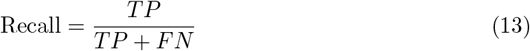

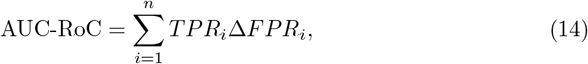

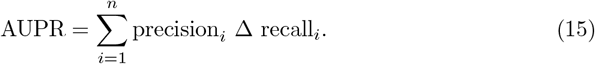

For imbalanced datasets, the Matthew Correlation Coefficient (MCC) is a robust metric that considers all values in the confusion matrix. MCC, unlike the F1-score [54], achieves high values only when all four confusion matrix parameters are high. We also prioritize the Area Under the Precision-Recall Curve (AUPR) due to the dataset imbalance. AUPR, which analyzes the precision-recall trade-off across different thresholds, provides a more informative evaluation than AUC-ROC [55].

As described in Section Basic definitions and methods, we outlined the various configurations used for Support Vector Machine (SVM) and Random Forest (RF). The best-performing SVM utilized a polynomial kernel with a C value of 10, while RF employed log loss as the criterion and set the maximum number of features to log *n*. The learning rate was 1 *×* 10^−5^. Additionally, we implemented 10-fold stratified cross-validation.

Performance was evaluated using four positive to negative (P2N) sampling ratios: 1:3, 1:5, 1:10, and 1:100. For example, a P2N ratio of 1:10, indicates that ten randomly selected negative samples were included for each positive sample.

Tables 2 through 5 present the results for the 1:3, 1:5, 1:10, and 1:100 P2N scenarios, respectively, reporting accuracy(Acc), MCC, AUC-ROC, and AUPR. Specifically, Table 2 shows results for the 1:3 P2N scenario. DRaW [18] (a CNN model) exhibited the lowest performance in this scenario. While other methods achieved near-perfect AUC-ROC and AUPR, Random Forest yielded the best accuracy. VCTatMLP and VCTatDot also demonstrated strong performance.

**Table 2.** Performance comparison for positive to negative sample ratio 1:3

| Method | Acc | MCC | AUC-RoC | AUPR |
| --- | --- | --- | --- | --- |
| SVM | $0.98 \pm 0.0$ | $0.96 \pm 0.0$ | **$0.99 \pm 0.0$** | **$0.99 \pm 0.0$** |
| RF | **$0.99 \pm 0.0$** | **$0.98 \pm 0.02$** | **$0.99 \pm 0.0$** | **$0.99 \pm 0.0$** |
| DRaW [18] | $0.94 \pm 0.04$ | $0.83 \pm 0.16$ | $0.97 \pm 0.02$ | $0.95 \pm 0.03$ |
| VCTatDot | $0.98 \pm 0.01$ | $0.95 \pm 0.02$ | **$0.99 \pm 0.0$** | **$0.99 \pm 0.0$** |
| VCTatMLP | $0.98 \pm 0.01$ | $0.96 \pm 0.02$ | **$0.99 \pm 0.0$** | **$0.99 \pm 0.0$** |

Table 3 presents the results for the P2N 1:5. A similar pattern is observed for performance; DRaW shows the lowest performance, while the attention-based methods demonstrate high performance. However, the random forest surpasses all methods in terms of MCC.

**Table 3.**
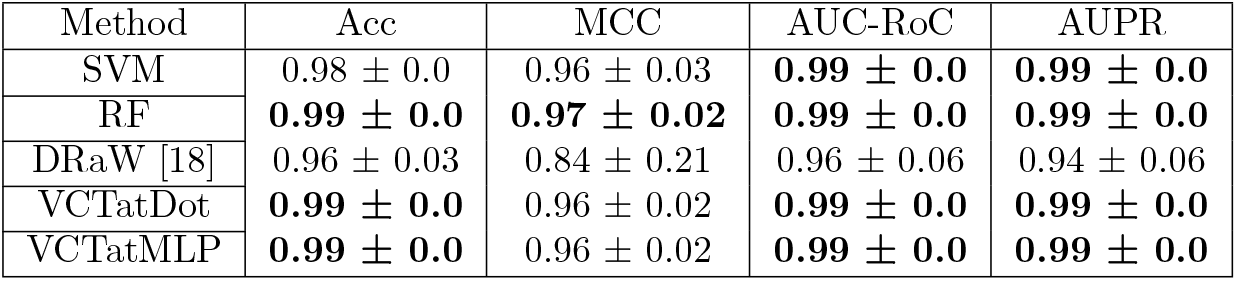
Performance comparison for positive to negative sample ratio 1:5

| Method | Acc | MCC | AUC-RoC | AUPR |
| --- | --- | --- | --- | --- |
| SVM | $0.98 \pm 0.0$ | $0.96 \pm 0.03$ | **$0.99 \pm 0.0$** | **$0.99 \pm 0.0$** |
| RF | **$0.99 \pm 0.0$** | **$0.97 \pm 0.02$** | **$0.99 \pm 0.0$** | **$0.99 \pm 0.0$** |
| DRaW [18] | $0.96 \pm 0.03$ | $0.84 \pm 0.21$ | $0.96 \pm 0.06$ | $0.94 \pm 0.06$ |
| VCTatDot | **$0.99 \pm 0.0$** | $0.96 \pm 0.02$ | **$0.99 \pm 0.0$** | **$0.99 \pm 0.0$** |
| VCTatMLP | **$0.99 \pm 0.0$** | $0.96 \pm 0.02$ | **$0.99 \pm 0.0$** | **$0.99 \pm 0.0$** |

Table 4 still indicates that DRaW has the poorest performance. Random forest surpasses other methods in terms of MCC and AUPR. VCTatDot and VCTatMLP demonstrate high performance as well.

**Table 4.** Performance comparison for positive to negative sample ratio 1:10

| Method | Acc | MCC | AUC-RoC | AUPR |
| --- | --- | --- | --- | --- |
| SVM | **$0.99 \pm 0.0$** | $0.96 \pm 0.02$ | **$0.99 \pm 0.0$** | $0.98 \pm 0.01$ |
| RF | **$0.99 \pm 0.0$** | **$0.97 \pm 0.02$** | **$0.99 \pm 0.0$** | **$0.99 \pm 0.0$** |
| DRaW [18] | $0.98 \pm 0.02$ | $0.84 \pm 0.25$ | $0.97 \pm 0.02$ | $0.93 \pm 0.04$ |
| VCTatDot | $0.98 \pm 0.0$ | $0.91 \pm 0.02$ | **$0.99 \pm 0.0$** | $0.98 \pm 0.0$ |
| VCTatMLP | **$0.99 \pm 0.0$** | $0.95 \pm 0.03$ | **$0.99 \pm 0.0$** | $0.97 \pm 0.01$ |

Table 5 shows that with the most challenging 1:100 positive-to-negative sampling ratio, VCTatDot outperforms other methods regarding three metrics: ACC, AUCROC, and AUPR. For the first time, it outperforms random forest regarding AUCROC. SVM has a similar performance to VCTatDot. Furthermore, VCTatDot outperforms VCTatMLP. Random forest performs better compared to other approaches regarding MCC.

**Table 5.**
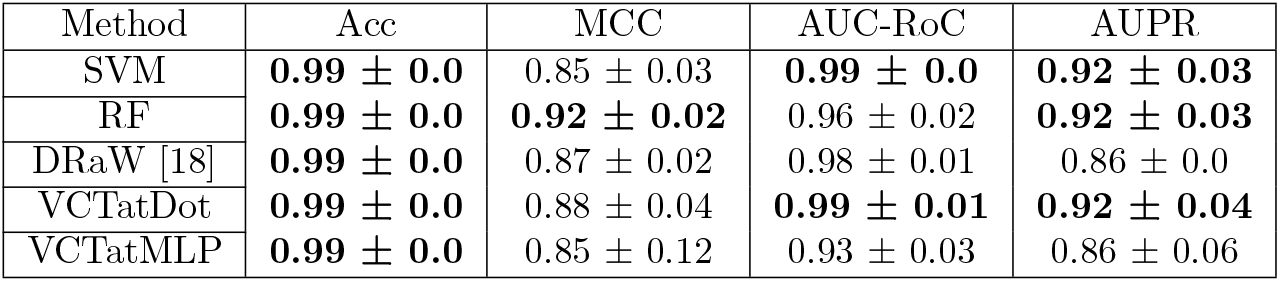
Performance comparison for positive to negative sample ratio 1:100

Figure 4 illustrates the loss convergence of VCTatMLP and VCTatDot, confirming the superior performance of VCTatDOT. The loss of VCTatMLP loss converges to 0.70, while VCTatDot’s loss converges to below 0.01, signifying an enhanced prediction of original labels due to the lower loss value. Furthermore, this improved convergence is observed in VCTatDot, which is a lighter and faster model than VCTatMLP.

**Fig 4.**
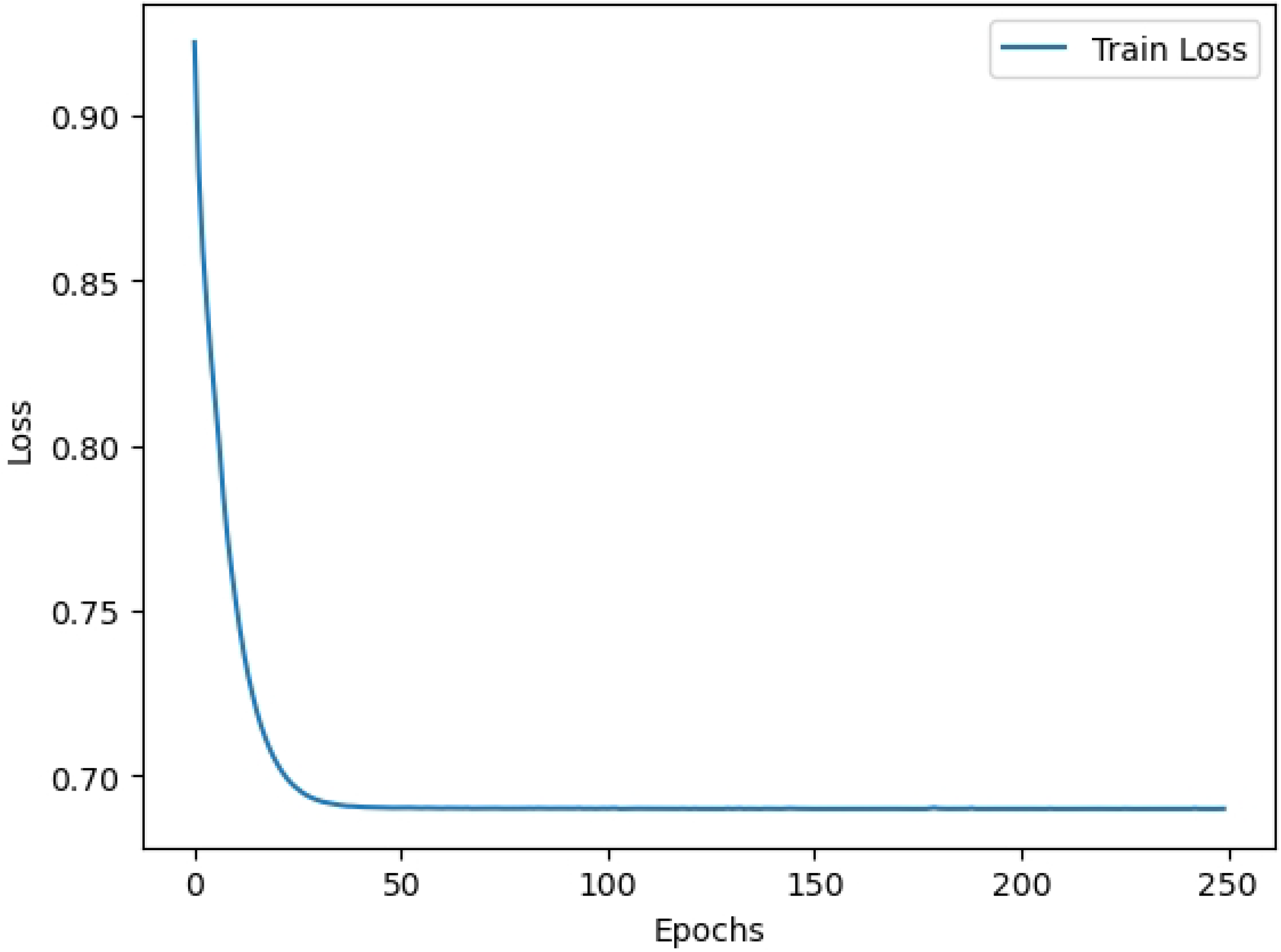
Convergence of loss functions for a positive to negative sampling ratio of 1:100. **(a)** VCTatDot **(b)** VCTatMLP

Figure 5 presents a schematic of the novel drug-drug-virus combinations predicted in this paper. Blue nodes represent drug combinations, and red nodes represent viruses. Edges indicate potential synergistic effects of the drug combination on the virus, with edge thickness reflecting the frequency of observed co-occurrences. For instance, the acyclovir and brincidofovir combination was observed 17 times in the test results.

**Fig 5.**
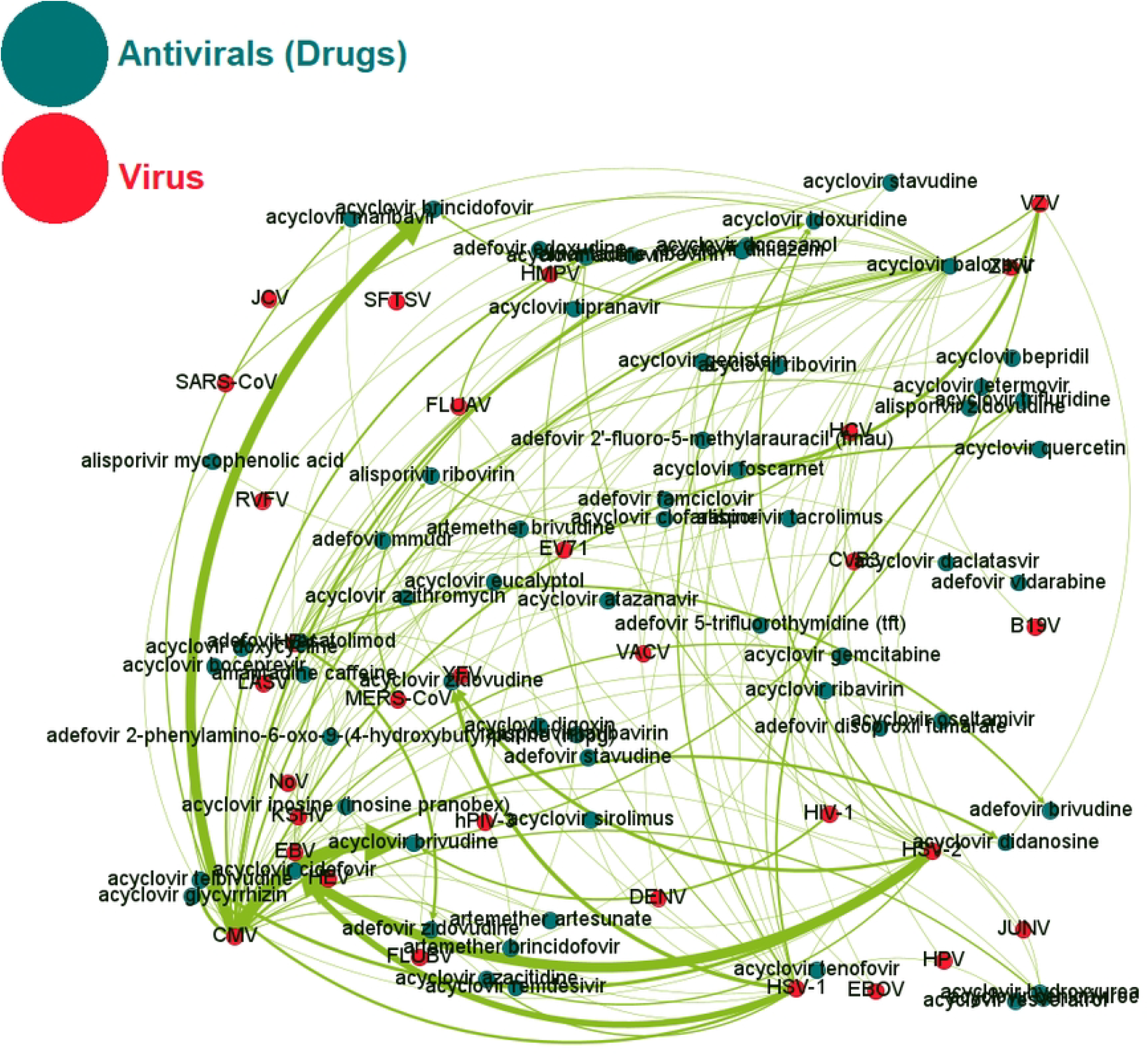
Predicted Drug Combinations. The blue vertices represent the combined drugs, while the red vertices indicate the viruses. Each edge suggests a synergistic effect of a drug combination on a virus.

Table 6 lists predicted synergistic drug pairs for viral diseases, focusing on combinations where each drug is independently prescribed for the corresponding virus and. For example, Acyclovir and Cidofovir are both prescribed for HSV-1. Critically, this work identified two synergistic combinations for HSV-1: acyclovir and ribavirin [56] and acyclovir and inosine [57]. Our methods correctly predicted both of these combinations.

**Table 6.** Predicted drug combinations that are either individually approved or shown to have synergistic effects in the literature.

| Antiviral 1 | Antiviral 2 | Virus |
| --- | --- | --- |
| Acyclovir [58] | Ribavirin [59] | HSV-1 [56] |
| Acyclovir [58] | Inosine (Inosine Pranobex) [60] | HSV-1 [57] |
| Acyclovir [58] | Cidofovir [61] | HSV-1 |
| Acyclovir [58] | Zidovudine [62] | HSV-1 |
| Acyclovir [58] | Didanosine [62] | HSV-1 |
| Acyclovir [58] | Genistein [63] | HSV-1 |
| Acyclovir [58] | Idoxuridine [64] | HSV-1 |
| Acyclovir [58] | Resveratrol [65] | HSV-1 |
| Acyclovir [58] | Stavudine [62] | HSV-1 |
| Acyclovir [58] | Adefovir [66] | HSV-1 |
| Acyclovir [58] | Trifluridine [67] | HSV-1 |
| Acyclovir [58] | Trifluridine [67] | HSV-2 |
| Acyclovir [68] | Cidofovir [61] | HSV-2 |
| Acyclovir [58] | Inosine (Inosine Pranobex) [60] | HSV-2 |
| Acyclovir [69] | Foscarnet [70] | VZV |
| Acyclovir [69] | Brincidofovir [71] | VZV |
| Acyclovir [69] | Brivudine [72] | VZV |
| Acyclovir [69] | Brincidofovir [71] | VZV |
| Alisporivir [73] | Ribavirin [74] | HCV |
| Alisporivir [73] | Mycophenolic Acid [75] | HCV |
| Alisporivir [73] | Taribavirin [76] | HCV |
| Acyclovir [69] | Maribavir [77] | CMV |
| Acyclovir [69] | Brincidofovir [78] | CMV |
| Acyclovir [79] | Foscarnet [70] | EBV |
| Amantadine [80] | Caffeine [81] | FLUAV |

## Conclusion

Synergistic combination therapy is valuable in drug repurposing for enhanced efficacy and reduced side effects. While computational drug discovery is rich with research in this area, studies on combination therapy for viral diseases remain scarce. This work introduces two attention-based approaches for synergistic antiviral combinations: one using an MLP, and the other a customized, tri-dot product. The tri-dot product, leveraging significant embeddings from attention-based generation, outperforms the MLP method, suggesting that substantial embeddings paired with a lighter prediction model are more effective. Also, our methods predict two drug combinations for the HSV-1 virus: acyclovir combined with ribavirin and acyclovir with inosine, both of which are approved by the literature. Furthermore, the proposed approach maintains and even improves its performance with increasing dataset size compared to other methods. These machine learning approaches also achieve high performance with reduced computational resources. Future work will focus on enhancing these attention-based approaches on larger datasets to further advance combination therapy for viral diseases. Additionally, XAI and explainable machine learning models are essential research in the files of predicting synergistic drug combinations for viruses.

## Data Availability

The datasets and code generated in this study are available in the GitHub repository: https://github.com/BioinformaticsIASBS/VirCombAtt.

## Author contributions statement

M.H. and S.M. conceptualized the study. M.H. provided supervision, managed the project, and led the writing of the original draft and subsequent revisions. S.M. contributed to method development, data curation, validation, and visualization, as well as the review and refinement of the manuscript.

---

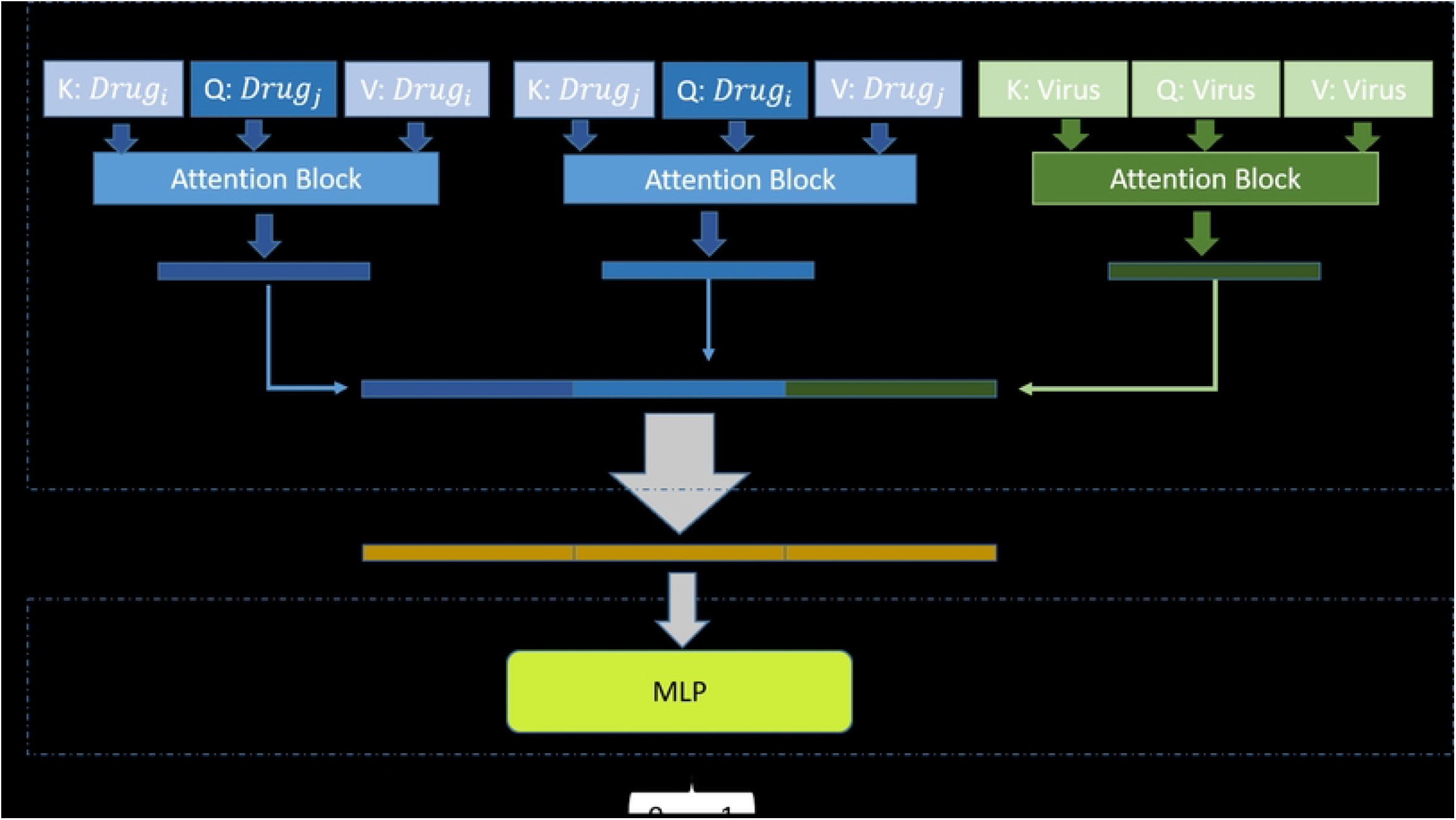

